# Processing Ergativity in Compound Light Verb Constructions: Electrophysiological Evidence from Hindi

**DOI:** 10.1101/2025.03.13.643010

**Authors:** Anna Merin Mathew, R. Muralikrishnan, Mahima Gulati, Shikha Bhattamishra, Kamal Kumar Choudhary

## Abstract

Ergativity marks subject arguments as agents of a transitive event and thereby signals verbal transitivity and influences language comprehension. We report here on an event-related brain potentials (ERP) study in Hindi, in which we investigated this interconnection to ascertain whether the ergative case as a processing cue and its ERP correlates can be generalized across and within ergative languages. The case marking on the subject argument (ergative or nominative case) in our study either matched or mismatched with the transitivity of the light verb (transitive or intransitive) in compound light verb constructions. Ergative case violations due to an intransitive light verb evoked an N400 effect, whereas nominative case violations due to a transitive light verb elicited a P600 effect. The results reveal neurophysiological differences in the processing of ergative and nominative case alignment modulated by the transitivity of the light verbs. The findings highlight the need for cross-linguistic research to aim beyond universality and elucidate the mechanism underlying the processing of language-specific structural variations.

## Introduction

Recent cross-linguistic research in the domain of sentence processing has aimed to identify the existing typological variabilities within languages and empirically examine their correlates at the neurobiological level (Bickel, Witzlack-Makarevich, Choudhary, Schlesewsky, and Bornkessel-Schlesewsky 2015; Bornkessel-Schlesewsky and Schlesewsky, 2016, 2019). This quest began with the recognition that in order to make a universal claim regarding the language comprehension mechanism, our understanding needs to be informed by empirical evidence from typologically diverse languages. However, this has been a challenge given that there exist many genetically and geographically disparate languages, with only a tiny proportion of them having been experimentally researched (Norcliffe, Harris, and Jaeger 2015). Among the many linguistic phenomena in the languages of the world, one well-known grammatical feature is ergativity (Comrie, 1978; Dixon, 1979; Delancy, 1981, 2006; Aldridge, 2008; Legate, 2012; Bhatt, 2007; Polinsky, 2007, 2016). The ergative case alignment pattern, found in about one-fourth of the world’s languages, has been widely studied from a theoretical perspective (Silverstein, 1976; Anderson, 1977; Comrie, 1978, 1981; Dixon, 1979, 1994; Foley and Van Valin, 1984; Woolford, 1997, 2015; Aldridge, 2008; Legate, 2008; McGregor, 2010; Verbeke, 2013, among many others). In many languages, case markers influence the morphological framework of words, the syntactic organization of the sentential structures, and the semantic composition of the entire sentence (Primus, 1999; Blake, 2001). Therefore, a pre-determined case marking system can alter how sentence structures are processed by the brain. In ergative languages, the agentive subject argument of transitive events is marked ergative, whereas the subject argument of intransitive events and the object argument of transitive arguments are unmarked (or identically marked). By contrast, in the more familiar accusative case alignment pattern, subjects of both transitive and intransitive constructions are unmarked (nominative), whereas object arguments are accusative case marked. The so-called split-ergative languages, such as Hindi (Mohanan, 1994), Tagalog (Aldridge, 2012), etc., show a split pattern, whereby they follow either the ergative or the accusative case alignment pattern depending on whether the sentence structure is transitive or intransitive. For example, in Hindi, when the verb is in the transitive form and in perfective aspect, the subject of the sentence is marked with an ergative case, whereas it is nominative in any other scenario (refer to section 1.2 for more details).

While several previous experimental studies have investigated the processing of case (Münte, Heinze and Mangun, 1993; Rösler, Friederici, Pütz and Hahne, 1993; Heinze and Münte, 1993; Münte, Matzke, and Johannes, 1997; Coulson, King, and Kutas, 1998; Wolff, Schlesewsky, Hirotani, Bornkessel-Schlesewsky, 2008; Frisch and Schlesewsky, 2001, 2005; Mueller, Hahne, Fujii, and Friederici, 2005, Mueller, Hirotani, and Friederici, 2007, among many others), just a few of them have examined the role played by the ergative case in sentence comprehension. An interesting question to consider in relation to the processing of ergative case is the extent to which the predictive ability of the ergative case marker differs or resembles that of other case markers and, additionally, whether the processing of the ergative case is influenced by structural variation. In other words, it is crucial to determine whether the processing of ergative case elicits similar or different neurophysiological signatures across ergative languages, as well as across different sentential structures within an ergative language. ERP studies exploring the processing of ergative case in ergative languages have probed the assumption of universality that underlines many psycholinguistic theories. The neurophysiological evidence from split-ergative languages such as Hindi (Choudhary, Schlesewsky, Roehm, and Bornkessel-Schlesewsky, 2009; Choudhary, 2011; Dillon, Nevins, Austin, and Phillips, 2012), Punjabi (Gulati, 2021) and ergative-absolutive languages such as Basque (Zawiszewski, 2007; Zawiszewski, Gutiérrez, Fernández, and Laka, 2011; Díaz, Sebastian Galles, Erdocia, Mueller, and Laka, 2011) suggest that ergative case indeed aids in the prediction of a transitive event similar to that of the accusative case in nominative-accusative case alignment languages. Moreover, the subject/agent preference appears to be a universal processing strategy applicable to ergative languages as well (Bickel et al., 2015). Although the aforementioned findings help generalize the processing mechanism, there is variability across the ergative languages in the neurophysiological correlates observed by these studies, indicative of processing differences between ergative and other types of case alignment systems. Taking a closer look at these existing studies (section 1.1) would reveal that the ERP effects elicited for the processing of ergative case varies not only across ergative languages (Hindi: Choudhary et al., 2009: N400-P600; Dillon et al., 2012: RAN-P600, Punjabi: Gulati, 2021: Late positivity) but also within a given ergative language (For instance, in Basque, Zawiszewski et al., 2011 reported a biphasic N400-P600 effect, whereas Díaz et al., 2012 observed a P600).

Furthermore, most of the aforesaid studies on the processing of ergative case have examined transitive sentences consisting of a simple predicate structure (i.e., marked by the presence of a single transitive verb). However, sometimes, it is not easy to define truly transitive or intransitive clause structure because that might be restricted to a particular feature and/or structure. It is for instance unclear whether the predictions held by an ergative case marker, or the interpretation of an ergative case marked sentence structure could change if placed within a complex predicate structure, with more than one verb symbolizing the event structure. The Hindi language is an ideal testing ground for investigating this as it is a split-ergative language containing multiple types of complex predicate structures, in which the subject case is determined by the transitivity of complex verbal predicates.

The split-ergativity system in Hindi is conditioned by the grammatical aspect and tense information on the verb (Dixon, 1979, 1994; Hale and Keyser, 1993; Piepers, 2016). Hindi exhibits morphological ergativity, whereby ergative structures are indicated through morphological case markers rather than syntactic arrangements (Woolford, 1997; Butt and Deo, 2001; Bhatt, 2007; Polinsky, 2007). The subject argument is marked with the ergative case only when its argument is agentive in nature, and when the verb is in the transitive form (active voice) marked with a perfective aspect. In such a construction, the direct object may be marked accusative or remain unmarked (nominative) depending upon the animacy of the object argument. Nevertheless, in all other aspectual forms, such as habitual or progressive, etc., the subject argument would be the unmarked nominative case. Further, the subject of an intransitive clause is never marked with an ergative case and remains unmarked by default (i.e., in the nominative case) regardless of the verbal tense/aspect. That is, the ergative case can only be assigned to truly transitive sentence structures (Kachru, 2006; Agnihotri, 2007). Whilst this is the general rule in Hindi, with a strong connection between ergativity and transitivity, there are certain exceptions. Thus, some unergative verbs can have ergative case-marked subjects, suggesting that the type of the verb can influence the split-ergativity pattern in Hindi (Dixon, 1979).

In this regard, a specific type of complex predicate structure in Hindi, the compound light verb structure^1^, contradicts the usual role of transitivity by offering an opportunity to examine the relationship between the case marking of arguments and the semantic as well as syntactic transitivity of verbs. Consider the following sentences (1a) and (1b), both denoting a similar event. Sentence (1a) is a simple predicate structure, whereas (1b) is a complex predicate structure known as the compound light verb construction in Hindi.

(1a.) *Mohan-ne aam khaa-yaa hai* Mohan.3SG.M-Erg mango.3SG.M.NOM eat-PFV.3SG.M Aux.SG.Prst ‘Mohan has eaten a mango’
(1b.) *Mohan-ne aam khaa li-yaa hai* Mohan.3SG.M-Erg mango.3SG.M.NOM eat take-PFV.3SG.M Aux.SG.Prst ‘Mohan has eaten a mango’

In (1a), a single verb conveys morphosyntactic as well as semantic information, including the transitivity of the sentence. The main verb *khaayaa* marks person, number, and gender agreement, shows the tense/aspect information and expresses the central meaning of the action of eating a mango in the sentence. The verb confirms the syntactic transitivity of the sentence since it assigns an ergative case to its arguments, and also the semantic transitivity of the sentence, which determines the number of arguments.

Sentence (1b) is an example of a compound light verb construction formed by the joining of two verbs acting as a single unit with their arguments mapped onto a monoclausal syntactic structure (Butt 1993a; 1995). Such a structure creates a bifurcation of the roles handled by a single verb in a simple predicate sentence onto two different verbs in a complex predicate structure. It is necessary to understand the roles these verbs perform to interpret the syntactic and semantic structure as well as to note how the light verb mediates the entire complex predicate structure despite being semantically bleached in itself.

In the compound light verb construction (1b), the first verb ‘*khaa*’ (meaning ‘eat’), which is in root/stem form (mainly in the non-inflected form, except in the instance of permissive light verb constructions), functions as a ‘polar verb’ (V1). The V1 retains its meaning such that it is replaceable with that of the whole compound light verb construction. The compound light verb structure ‘*khaa liyaa*’ conveys ‘the act of having eaten a mango’, deriving its predominant semantics from V1. Furthermore, the transitivity of V1 determines the event structure of the whole light verb construction, i.e., it determines the number of arguments mandatorily required. An essential thing to note here is that V1 is never inflected for its grammatical relation with the subject or object in the sentence. Rather, the second verb ‘*liyaa*’ (meaning ‘take’), functions as an ‘explicator verb’ (V2) (Abbi and Gopalakrishnan, 1991) and is also known as the ‘light verb’ (Jespersen, 1965), vector verb (Dasgupta, 1977), intensifying verb, operator verb, or compound auxiliary. (Butt, 1995). It is said to be semantically bleached since V2 contributes relatively less towards the meaning of the compound light verb structure. However, unlike V1, the light verb V2 not only assigns the case marking of the arguments, but also shows person, number, and gender agreement, and is marked for tense, modal, and aspectual information. It also expresses certain extra-linguistic information such as contributing a sense of perfectivity (Hook, 1974), forcefulness, suddenness, volitionality (Mohanan, 1994), agentivity, benefaction, and direction (Hook, 1974; Mohanan, 1994), as well as enhances properties such as finality, definiteness, manner of the action, attitude/intention of the speaker, a sense of negative value, etc. (Hook, 1974; Butt and Geuder, 2001; Butt, 1993b; Abbi, 1992; Abbi and Gopalakrishnan, 1991). For this reason, even though the light verb V2 is referred to as ‘light’ and is considered to be semantically bleached, it is not empty in any sense. Rather, it is of immense importance as it is responsible for case assignment. It is the ± transitivity of the light verb that determines the syntactic (structural) transitivity of the compound verb construction (Masica, 1976; Abbi and Gopalakrishnan, 1991). In example (1b), it is the transitive and perfective light verb *liyaa* (meaning ‘take’) that assigns ergative case to the subject argument, not the transitive main verb *khaa* (meaning ‘eat’). The (in)transitivity of light verbs plays a major role in governing ergativity in Indo-Aryan languages (Amritavalli, 1979; Bhatt, 2007; Mahajan, 2012).

On the other hand, some perfective sentences in Hindi occur within the compound light verb structure pattern, but cannot assign an ergative case to their subject argument. This happens in a specific kind of compound light verb construction, in which there is a change in the transitivity of verbs within the complex predicate. Consider the sentence in (1c).

(1c.) *Mohan aam khaa ga-yaa hai* Mohan.3SG.M.NOM mango.3SG.M.NOM eat go-PFV.3SG.M Aux.SG.Prst ‘Mohan has eaten a mango’

In (1c), there are multiple markers of transitivity. There is a human animate argument marked with ergative case (indicative of being a prototypical agent) and the presence of a prototypical patient argument (low on animacy scale), i.e., an inanimate object and the affected one, by virtue of which the animate argument is reiterated as the subject (Hopper and Thompson, 1980; Dowty, 1991; de Hoop and Narasimhan, 2005). Here, the polar verb V1 is transitive and capable of assigning two thematic roles, namely actor and undergoer, because of which the sentence is semantically transitive. Therefore, the polar verb V1 determines the number of arguments compulsorily required in the sentence (two roles in (1c)), and thus also determines the event structure. However, it still cannot assign case to the actor/subject argument. The light verb V2 in (1c) is intransitive *gayaa* and hence cannot assign an ergative case to the actor/subject argument. Consequently, the subject here is in the default nominative case. Syntactically, the transitivity of the light verb is reduced because an intransitive light verb can no longer mark an ergative case to the agent argument (*Mohan*) of the sentence (Nespital, 1997; Drocco, 2018; Drocco and Tiwari, 2020). This demonstrates how case assignments are solely based on the transitivity of the light verb in such compound light verb constructions, which adhere to a specific split-ergative rule combining the features of ergativity and transitivity intricately.

On the basis of the relationship between the semantic and morpho-syntactic transitivity, and grounded on the differences between meaning and structure, it is often challenging to call only one of the above-discussed constructions (1b and 1c) a prototypical transitive sentence when compared with the simple transitive sentence as in (1a). An attribute of the compound light verb construction is that although both the verbs V1 and V2 have separate functions, these verbs nevertheless together form a verbal complex that fulfils the fundamental characteristics of complex predication, and function grammatically and semantically as a single entity. In the above-discussed examples, the verbal predicate structures, *khaayaa, khaa liyaa*, *khaa gayaa,* all denote an act of ‘having eaten’. In spite of that, they are quite different in the way they express semantic and syntactic information.

Given this scenario, we investigate in the present study, the neural correlates of ergative case processing in compound light verb constructions to examine whether the underlying mechanism behind the computation of the ergative case-marked construction is similar to that of the default nominative case-marked construction. We also examine whether the findings are comparable to that of existing studies from ergative languages. Finally, we also inquire into the symbiotic relationship between the alternating case marking pattern of the subject argument and the changing degree of verbal transitivity in complex verbal predicate structures during real-time language comprehension. The following paragraphs describe the previous electrophysiological findings from ergative languages in detail (section 1.1), before moving on to the present study (section 1.2).

### Electrophysiology of Ergative Case

Several studies have inquired into the processing of case to understand the mechanisms underlying different case alignments and their prediction regarding language comprehension. These have examined a number of languages to understand how crucial sentential information -- such as the construction of core arguments and thematic roles as well as their relation to the verb/verbal phrase, -- are determined and encoded within specific linguistic features that act as processing cues. Case markers are one of the syntactic encoders (among other syntactic and semantic information types like agreement, animacy, word order, etc.) of sentential information. Case markers distinguish sentential arguments, play an important role in the assignment of actor and undergoer roles, and function as linguistic cues that aid in sentence processing. In addition, case markers help in identifying ambiguous arguments (often non-case marked and in the first position), which are frequently identified/preferred as subjects/agents (Dutch: Frazier and Flores d’Arcais, 1989; Italian: de Vincenzi, 1991; German: Bornkessel, McElree, Schlesewsky, & Friederici 2004, Haupt, Schlesewsky, Roehm, Friederici Bornkessel-Schlesewsky, 2008; Turkish: Demiral, Schlesewsky, Bornkessel-Schlesewsky, 2008, Chinese: Wang, Schlesewsky, Bickel, & Bornkessel-Schlesewsky, 2009; Swedish: Hörberg, Koptjevskaja-Tamm, and Kallioinen, 2013; Hindi: Choudhary et al., 2009, 2011; Bickel et al., 2015). They also help form a dependency relationship between constituents, enabling the prediction of upcoming arguments with specific features such as animacy, specificity/definiteness, etc. Therefore, case markers ensure the establishment of thematic hierarchization between the arguments and, if necessary, can signal the need for thematic reanalysis as well (German: Bornkessel, Schlesewsky, Friederici, 2003, Roehm, Schlesewsky, Bornkessel, Frisch, Haider, 2004, Bornkessel, Fiebach, Friederici, 2004, Frisch and Schlesewsky, 2001, 2005; Japanese: Wolff, Schlesewsky, Hirotani, Bornkessel-Schlesewsky, 2008; Chinese: Philipp, Bornkessel-Schlesewsky, Bisang, & Schlesewsky, 2008; Spanish: Nieuwland, Martin, and Carreiras, 2013; Hindi: Schlesewsky, Choudhary, and Bornkessel-Schlesewsky, 2010). Moreover, case markers at the argument position itself can build an expectation of the verb and its category as well as predict features associated with the verb, such as tense, aspect, and agreement information like gender, number, and person (Icelandic: Bornkessel-Schlesewsky, Roehm, Mailhammer, Schlesewsky, 2020; Hindi: Nevin et al., 2007, Choudhary et al., 2009, 2011; Bickel et al., 2015). Additionally, case markers also give rise to predictions about the upcoming sentence structure and possible interpretations (Japanese: Wolff et al., 2008; Hindi: Choudhary et al., 2009, Sauppe, Choudhary, Giroud, Blasi, Norcliffe, Bhattamishra, Gulati, Egurtzegi, Bornkessel-Schlesewsky, Meyer, Bickel, 2021).

Further, they provide vital information for facilitating processing strategies for different kinds of structural, thematic and grammatical function reanalyses (Friederici, 1995, 2002; Friederici and Mecklinger, 1996; Friederici, Mecklinger, Spencer, Steinhauer, and Donchin, 2001; Bornkessel, McElree, Schlesewsky, Friederici, 2004; Frisch, Schlesewsky, Saddy, Alpermann, 2002; Schlesewsky & Bornkessel, 2006). Findings from behavioural and ERP studies from the language production domain reveal that the case morphology of a language can shape the relational, structural and linguistic encoding processes. Case can affect the time of verb retrieval and help anticipate the verb-final structure (Hindi: Husain, Vasishth, Srinivasan, 2014; Zafar and Husain, 2022), the early stage of sentence planning (Hindi: Sauppe et al., 2021; Basque and Swiss German: Egurtzegi, Blasi, Bornkessel-Schlesewsky, Laka, Meyer, Bickel, Sauppe, 2022), and especially assist in visual apprehension of events (Basque and Spanish: Isasi-Isasmendi, Sauppe, Andrews, Laka, Meyer, and Bickel, 2023; Norcliffe et al., 2015; Tzeltal and Tagalog: Sauppe, Norcliffe, Konopka, Valin, and Levinson, 2013, Russian and Hebrew: Meir, Parshina, Sekerina, 2020).

In view of these findings, primarily drawn from research on nominative-accusative languages, a recent strand of research has been focusing on processing differences in ergative languages, by examining linguistic features in these languages that are quite different from that of nominative-accusative languages. Taking an experimental approach to address ergativity from a variety of directions (Longenbaugh and Polinsky, 2017; Zawiszewski, 2017, for an in-depth review of cross-linguistic evidence), this emerging body of research questions the universality of language processing strategies with respect to the processing of ergative case. In this context, some studies have investigated the processing of filler-gap dependencies to understand the relationship between case marking and grammatical function. Specifically, they tested whether an ergative and/or absolutive subject preference could be established over an object preference in ergative languages and also verified if ergative languages favour an ergative subject above any other kind (Basque: Carreiras, Duñabeitia, Vergara, de la Cruz-Pavía and Laka, 2010; Avar: Polinsky, Gallo, Graff and Kravtchenko, 2012; Ch’ol and Q’anjob’al: Clemens, Coon, Pedro, Morgan, Polinsky, Tandet, Wagers, 2015; Niuean: Longenbaugh and Polinsky, 2017). Carreiras et al., (2010) studied whether subject relative clauses (SRC) are easier to process than object relative clauses (ORC) in Basque, like in nominative-accusative languages (Friederici, Mecklinger, Spencer, Steinhauer and Donchin, 2001; King and Kutas, 1995; Weckerly and Kutas, 1999). They conducted two self-paced reading studies and an ERP study, in which the critical position at which both sentences disambiguated either to an SRC or ORC interpretation was at the sentence-final auxiliary verb. The results from their self-paced reading study showed that object-relative clauses took less time to read and were easier to process. Their ERP results revealed a larger P600 for subject-relative than object-relative clauses, indicating that SRCs are harder to process. The authors observed that in Basque, objects are not marked (default), but rather, transitive subjects are marked, and thus, object-relative clauses were easier to process.

Polinsky et al., (2012) used self-paced reading to test whether subject-relative clauses are universally more effortlessly processed than object-relative clauses by testing subject preference and ergativity in Avar and did not observe a processing difference between the ergative subject and the absolutive object in Avar. On the other hand, Bickel et al., (2015) examined subject/agent processing preference in an ERP study on Hindi to determine whether its universality extends to ergative languages as well. They employed fully grammatical sentences (OVS order) that varied in the sentence-initial inanimate argument, which was either unambiguous by virtue of being accusatively case-marked or ambiguous with no overt case marking. The verb that followed was either imperfective or in perfective aspect. They found that, in conditions with an ambiguous first argument, i.e., not case marked, the argument was initially processed as a subject/agent of the sentence; but on encountering the verb, the argument was reanalyzed as the object of the sentence. That is, the verb information disambiguated the initial ambiguous noun to be an undergoer. This thematic reanalysis engendered an N400-P600 effect at the verb, regardless of its aspect type. These studies reveal that language processing strategies differ across ergative languages.

Previous ERP studies on the processing of ergative case have attempted to establish the cross-linguistic validity and the extent to which the electrophysiological correlates elicited for processing case in nominative-accusative languages apply to ergative languages. In an ERP study in Hindi, Choudhary, Schlesewsky, Roehm, and Bornkessel-Schlesewsky (2009) investigated the processing of ergative and nominative subject cases to test whether the previously obtained ERP components for the processing of case (N400 in Frisch and Schlesewsky, 2001; Haupt et al., 2008) could be due to the reason that case belongs to a morphosyntactic category that brings about changes in the construal of the sentence as well. An auditory presentation of simple transitive sentences was employed, in which the subject case of the first argument, ergative or nominative, and the aspect of the transitive verb, perfective and imperfective, were manipulated. The critical position was the aspect marker on the simple transitive verb, which determines the grammaticality of the subject case in Hindi. For a sentence to be grammatical, the aspect marker on the verb must be obligatorily perfective for an ergative marked argument (non-default case), while it must be imperfective for a nominative case marked argument (default case). (2) Examples of the experimental stimuli from Choudhary et al. (2009).

a. *shikshak maalii-ko dekh-taa hai* teacher.NOM gardener-ACC see-IPFV.3SG.M AUX.PRS ‘The teacher sees the gardener.’
b. * *shikshak-ne maalii-ko dekh-taa hai* teacher-ERG gardener-ACC see-IPFV.3SG.M AUX.PRS ‘The teacher sees the gardener (intended).’
c. * *shikshak maalii-ko dekh-aa hai* teacher.NOM gardener-NOM see-PFV.3SG.M AUX.PRS ‘The teacher has seen the gardener (intended).’
d. *shikshak-ne maalii-ko dekh-aa hai* teacher-ERG gardener-ACC see-PFV.3SG.M AUX.PRS ‘The teacher has seen the gardener.’

Ergative case violations (2b) elicited a biphasic N400-P600 effect, whereas nominative case violations (2c) elicited an N400 effect compared to their respective non-anomalous counterparts. The N400 and P600 had a larger amplitude for ergative case violations. The N400 effect was attributed to an interpretively relevant rule mismatch, and its larger amplitude was said to be due to the mismatch being more problematic for the ergative violation conditions. The late positivity effect ensued only for ergative case violations because the ergative case in Hindi is a non-default rule that applies merely to one restricted environment, i.e., when the transitive verb is in the perfective aspect. Thus, an over-application of the non-default rule led to the ill-formedness of the construction and engendered a P600 effect, interpreted as a marker for well-formedness-related issues (Bornkessel & Schlesewsky, 2006) and conflict monitoring (van Herten et al., 2006).

Similar to the study by Choudhary et al., (2009), Gulati (2021) conducted a visual ERP study in Punjabi, a sister Indo-Aryan language that exhibits aspect and person-based split ergativity, to investigate whether the neural correlates of processing subject case violations in Punjabi would be similar to that of Hindi. This study also employed a mismatch between the case marking of the sentence-initial subject argument (ergative and nominative) and the aspect of the critical verb (imperfective and perfective). The findings, in contrast to results from Hindi, showed an early positivity (300-500ms) for the nominative case violation and a late positivity (500-700ms) for the ergative case violation. Gulati (2021) concluded that these ERP differences are indicative of the fact that the ergative case might not be as strong a cue in Punjabi for argument and structure interpretation, unlike in Hindi. This suggests that native speakers of Punjabi perhaps draw significant inferences from other linguistic cues such as word order etc., when processing these structures. These results from Hindi and Punjabi show that universality for the processing of ergative case cannot be established even within typologically similar languages, and that the grammatical properties specific to a given language are crucial.

Further evidence to this argument comes from Basque, an ergative language that is very different from Hindi and follows an ergative-absolutive case-marking system (Dixon, 1994; Marantz, 1984). Diáz, Sebastián-Gallés, Erdocia, Mueller, and Laka, (2011) conducted an auditory ERP study on Basque to investigate case and agreement processing, in which participants listened to grammatical sentences and ungrammatical sentences with double ergative case violations in the SOV word order. The authors aimed to compare the ERP correlates of processing such structures in Basque with the previously reported work that examined double case violations (double nominative, accusative and dative cases) in German, in which such violations elicited an N400-P600 effect (Frisch and Schlesewsky, 2001, 2005; Mueller, Hirotani & Friederici, 2007). When both the subject and object arguments were marked ergative in Basque, the second ergative case marked argument evoked a large positivity between 400 and 1250ms. This result, whilst similar to findings from German, differed from findings from Hindi, in which an N400 effect was evoked (Choudhary et al., 2009). The authors proposed that these ERP differences arose due to the underlying difference in the ergative alignment between Hindi and Basque. Hindi has a split-ergative pattern, in which the agent is assigned ergative case only in the perfective aspect (and mostly with transitive verbs). By contrast, Basque exhibits a truly ergative alignment whereby all agents are marked with an ergative case without exception. Sentences employed in the Basque experiment by Díaz et al., (2011) always had correctly (ergative) marked arguments, which did not probably cause semantic difficulty in interpreting the agency of the ergative marked arguments, contrary to the Hindi study. Furthermore, in Basque, some sentences with a second ergative case marked argument do not make the sentence completely ungrammatical because they can be temporarily considered as the subject of an embedded clause. Thus, there was no N400 effect found in the Basque study. As pointed out by Díaz et al., (2011), this variation is suggestive of the fact that even within ergative languages, the differences in the grammatical characteristics of the particular language would lead to very different results. Nevertheless, the P600 effect, which was found in both Hindi and Basque, appears to suggest that the detection of case violations is similar across these ergative languages and does not depend on the argument alignment type.

Zawiszewski, Fernández, Gutiérrez, and Laka (2011) examined how native Basque speakers who spoke Spanish as a second language and non-native Basque speakers who spoke Spanish as their first language processed the ergative case in Basque. Grammatical and ungrammatical OVS sentences were compared, with the order becoming clear only at the second argument that either was ergative (i.e., agent and thus non-anomalous OVS order), or was unmarked (i.e., anomalous, because of no agent argument). The ergative case violation elicited a biphasic N400-P600 pattern in native speakers of Basque, while evoking only an N400 effect in non-native speakers. The N400 was said to result from the difficulty in assigning a thematic role to the agent argument that lacked ergative case marking. The P600 effect was only evoked in native speakers, indicative of the fact that they had the ability to identify ill-formed sentences that the non-native speakers perhaps did not have. The authors speculated the possibility that the brains of non-native speakers of Basque perhaps used a processing strategy that was transferred from their first language, Spanish (a nominative-accusative language), which rendered the ergative case violations as relatively acceptable for them. This could be because the Spanish native speakers would/could interpret Basque sentences with an agent argument that lacks an ergative case simply as a nominative argument. According to Zawiszewski et al., (2011), non-native Basque speakers ignore ergativity and derive their subjecthood from other characteristics, such as animacy. Hence, there is no need for reanalysis or repair, thus resulting in the absence of the P600 effect.

Taking a slightly different approach, Dillon et al., (2012) focused on syntactic and semantic cues to language processing to understand how these predictors/cues affect tense processing in Hindi. The authors controlled the expectation about the verbal morphology in two ways: by using a past tense adverb, which requires a past tense marked verb (semantic cue), and employing an ergative case marked argument, which needs a verb in perfective aspect (syntactic cue). The critical verb was in the past tense (i.e., non-anomalous), or alternatively, it was in the future tense (i.e., a tense violation). Results at the verb showed different ERP responses depending on which type of cue predicted the verbal tense form. When the ergative case (syntactic cue) was predicting a verb type, and there was a tense violation at the verb, a right anterior negativity RAN effect and a P600 effect were elicited. In contrast, when the past tense adverb (semantic cue) predicted the verbal tense type, a violation of the verb tense elicited an early posterior negativity (200-400ms) and a P600 effect (600-800ms). While both the violations engendered a positivity effect, it was larger for the ergative case-based violation conditions, suggesting that the prediction for a past tense verb due to the ergative case on the first argument was stronger.

Taken together, the evidence from comprehension studies suggests that the relationship between ergative case and other linguistic features that affect the processing of case, such as transitivity, aspect, tense, and agreement, remains inconclusive. For instance, the ERP correlates obtained for ergative case violations differ depending upon the kind of structures involved, and on whether the manipulation was exclusively based on case assignment and/or with other intervening factors. Another factor to consider is that previous studies on Basque contrasted arguments that are ergative marked versus null marked, either at the argument position itself or at the disambiguating verb (Díaz et al., 2011 and Zawiszewski et al., 2011). Studies from Hindi and Punjabi violated the aspectual requirement of a split-ergative case alignment system, with the result that the ergative or nominative case violation would be realized on the violated perfective or imperfective aspect marker of the verb (Choudhary et al., 2009, 2011; Dillion et al., 2012; Gulati, 2021). In order to disentangle the effect of case from that of aspect or tense however, it would be necessary to elicit ergative structure violations without simultaneously violating other features of the sentence, such as aspect or tense. This would be possible in Hindi light verb constructions, in which the aspect/tense information can be kept constant while manipulating the transitivity of the light verb. Thus, Hindi light verb constructions allow determining whether and how a variation in the verbal transitivity of a complex predicate construction influences the case processing at the sentence-final verb position.

The processing and representation of light verb constructions have been previously examined through a multidisciplinary approach, combining behavioural and neurophysiological methods with insights from theoretical linguistics, including the separate entry and under-specification approaches (see Wittenberg and Piñango, 2011), and the language-cognition interface models (Hale and Keyser, 1993, 2002) and Parallel architecture framework (Culicover and Jackendoff, 2005). Some of these studies observed that light verb forms require more mental processing than non-light forms, resulting in higher processing costs and slower reaction times. The event co-compositional nature of light verb constructions was stated as a reason for a lack of canonical mapping between semantics, syntax, and cognitive event structures in real-time processing (English: Piñango, Mack, and Jackendoff, 2006; German: Wittenberg and Piñango, 2011; Wittenberg and Snedeker, 2014; Wittenberg, Khan, and Snedeker, 2017, while some studies reported contradictory results, in German (Briem, Balliel, Rockstroh, Butt, Schulte im Walde, and Assadollahi, 2010), and in Hindi (Vaidya and Wittenberg, 2020). Wittenberg et al., (2014) conducted an ERP study in German that compared light verb constructions (the pairing of a light verb with an eventive noun) with different non-light constructions, in which they observed a frontally focused late sustained negativity (500-900ms) for light verbs. Several studies in Hindi have used conjunct light verb constructions to study sentence processing, event understanding, and morpho-syntactic properties. Specifically, studies have investigated expectation and memory in sentence processing (Husain et al., 2014), perfective processing in event understanding (Arunachalam and Kothari, 2011), and agreement attraction and case-agreement interactions (Bhatia and Dillon, 2022). In Chinese, Liao, Chia-Hsuan, Lau, and Ellen (2020) were able to observe processing differences in the encoding of complex events by analyzing different kinds of compound verb constructions. They found an N400 when complex verbs predicted object nouns. To sum up, the majority of the research mentioned here illustrates a fundamental distinction between the neural processes governing simple and complex predicate structures. However, there is a relative paucity of conclusive findings about the processing of complex predicate structures, particularly about whether they show qualitatively similar neurophysiological correlates across languages.

### The Present Study

We present here a visual ERP study, in which we investigated how ergativity is processed in Hindi compound light verb constructions. We examine how the brain processes ergative and nominative case markers, and how they influence predictions about upcoming arguments, verbal features, and sentence structure. In other words, our aim is to elucidate whether initial case-based predictions are revised upon encountering complex predicate structures, and how the transitivity of light verbs interacts with the prediction based on subject case markers, particularly when case assignment is realized on the light verb itself.

In this regard, we designed a 2 x 2 experiment manipulating the morphological case of the subject argument (nominative case/ ergative case) and the transitivity of the light verb (transitive light verb/ intransitive light verb) in compound light verb predicate constructions. This resulted in four critical conditions as shown in Table 1: Ergative Transitive (ET), Nominative Intransitive (NI), Ergative Intransitive (*EI), and Nominative Transitive (*NT). All sentences were of the form Adverb – NP1– NP2 – V1 – V2 – Aux. NP1 was the animate subject and was either ergative case marked or was in the default nominative case; NP2 was the unmarked inanimate object; V1 was the polar verb in the transitive root form, devoid of grammatical markers; and V2 was the light verb whose transitivity either matched or violated the subject case assignment. The sentence-final auxiliary indicated present tense. The critical position was at the light verb in line with the previous ERP studies in Hindi (Choudhary et al., 2009; Choudhary, 2011; Dillon et al., 2012).

**Table 1:**
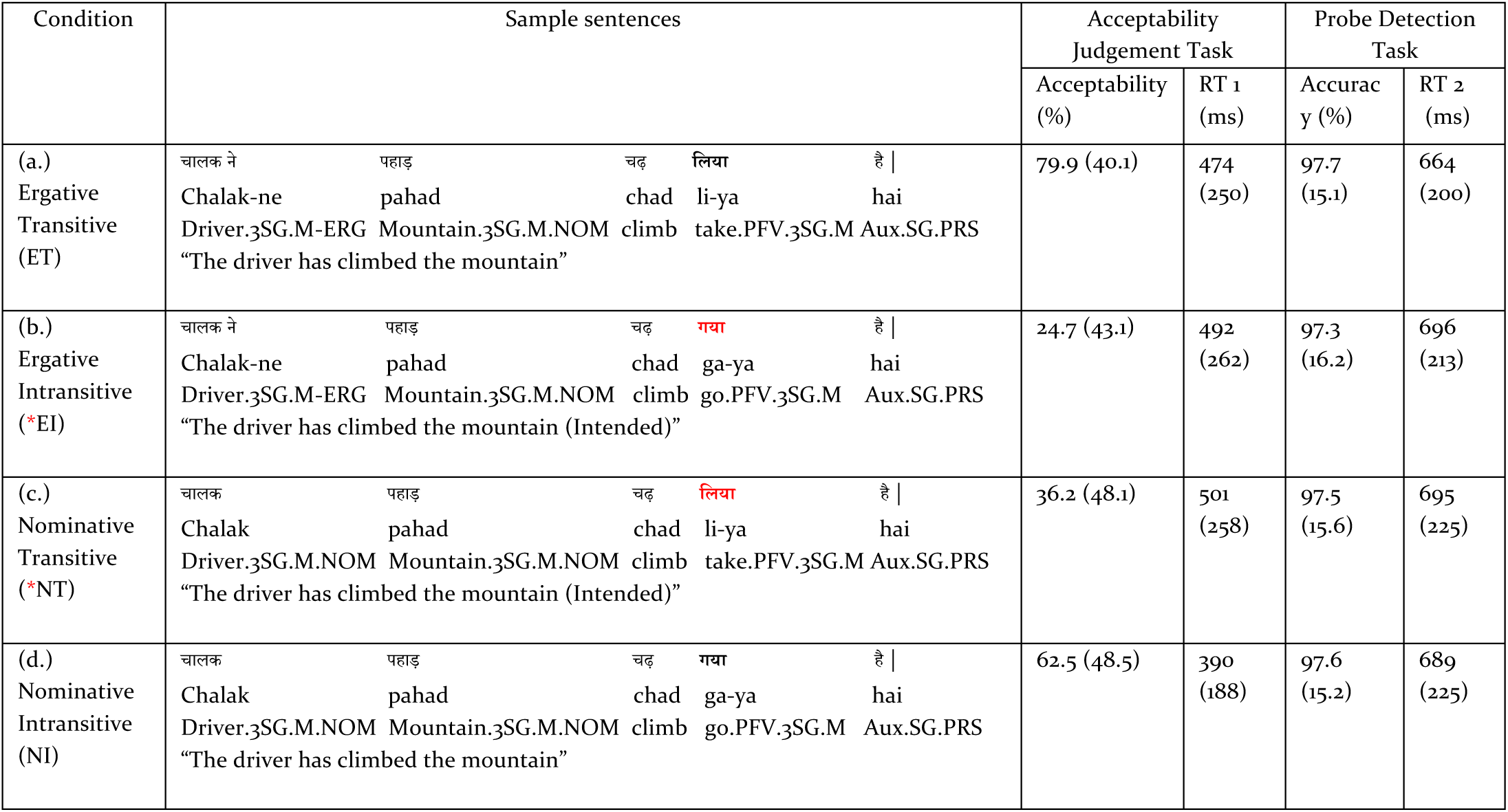
Experimental Sample The critical position is the light verb, which is marked in boldface. The use of an asterisk ‘*’ indicates a violation of the subject case marking rule, depicting a mismatch between the subject case and the transitivity of the light verb. The values represent the mean acceptability and mean accuracy (%) and the respective reaction times (ms) for the two behavioural tasks per four conditions, with the standard deviations provided in parentheses (values have been rounded off). Abbreviations: ERG: Ergative; NOM: Nominative; PFV: Perfective Aspect of the verb; M: Masculine; F: Feminine; SG: Singular; 3: Third Person, Aux: Auxiliary, PRS: Present tense. Acceptability: mean

| Condition | Sample sentences | Acceptability Judgement Task |  | Probe Detection Task |  |
| --- | --- | --- | --- | --- | --- |
|  |  | Acceptability (%) | RT 1 (ms) | Accuracy (%) | RT 2 (ms) |
| (a.)<br>Ergative Transitive (ET) | चालक ने पहाड़ चढ़ लिया है \|<br>Chalak-ne pahad chad li-ya hai<br>Driver.3SG.M-ERG Mountain.3SG.M.NOM climb take.PFV.3SG.M Aux.SG.PRS<br>“The driver has climbed the mountain” | 79.9 (40.1) | 474 (250) | 97.7 (15.1) | 664 (200) |
| (b.)<br>Ergative Intransitive (*EI) | चालक ने पहाड़ चढ़ गया है \|<br>Chalak-ne pahad chad ga-ya hai<br>Driver.3SG.M-ERG Mountain.3SG.M.NOM climb go.PFV.3SG.M Aux.SG.PRS<br>“The driver has climbed the mountain (Intended)” | 24.7 (43.1) | 492 (262) | 97.3 (16.2) | 696 (213) |
| (c.)<br>Nominative Transitive (*NT) | चालक पहाड़ चढ़ लिया है \|<br>Chalak pahad chad li-ya hai<br>Driver.3SG.M.NOM Mountain.3SG.M.NOM climb take.PFV.3SG.M Aux.SG.PRS<br>“The driver has climbed the mountain (Intended)” | 36.2 (48.1) | 501 (258) | 97.5 (15.6) | 695 (225) |
| (d.)<br>Nominative Intransitive (NI) | चालक पहाड़ चढ़ गया है \|<br>Chalak pahad chad ga-ya hai<br>Driver.3SG.M.NOM Mountain.3SG.M.NOM climb go.PFV.3SG.M Aux.SG.PRS<br>“The driver has climbed the mountain” | 62.5 (48.5) | 390 (188) | 97.6 (15.2) | 689 (225) |

In compound light verb constructions, the transitivity and perfectivity of the light verb determine the correct case marking on the subject argument (Amritavalli, 1979). Thus, the violation conditions in the present study do not constitute any agreement or aspect violations, but rather are entirely based on whether the light verb is transitive or not. In other words, at the position of the light, the two violation conditions (*EI and *NT) had a mismatch between the case marking of the subject and the transitivity of the light verb, which contradicted the subject case-based prediction for a transitive versus intransitive light verb. Any differences in the ERPs at this position in comparison to the non-anomalous counterparts would then be a reflection of the unmatched prediction for a transitive or intransitive light verb.

Our hypotheses were as follows. If the predictions that the two kinds of subject cases (nominative and ergative) give rise to are similar, and if nominative and ergative subjects are processed similarly, then the ERPs at the position of the light verbs should be qualitatively similar for nominative and ergative case violations. Alternatively, based on previous findings from Hindi (Choudhary et al., 2009), we hypothesize that processing differences between ergative and nominative case violations should emerge at the position of the light verbs, with ergative case violations evoking a biphasic N400-P600 effect and nominative case violations engendering an N400 effect. However, there is a crucial difference between the stimuli employed in previous Hindi studies and our study, namely the structural differences due to the use of compound light verb structures. That is, the assignment of the subject case in our stimuli is regulated by the transitivity of the light verb V2. Further, everything up until the light verb remains constant and there is no other morphosyntactic violation occurring at any other position. In view of this, if results obtained at the light verb are similar to previous findings, this would indicate that case-based violations are processed qualitatively similarly across diverse sentence structures in Hindi. By contrast, if the use of compound light verb structures engenders a different pattern of results compared to previous findings, this would suggest that the role that the ergative case plays in language comprehension, both within specific languages and across language-specific structures, cannot be generalized.

## Materials and Methods

### Participants

Thirty first-language speakers of Hindi (mean age: 20.5 years, range: 18-30), mostly graduate students and staff at the Indian Institute of Technology Ropar, India, participated in the study. They predominantly hailed from Hindi-speaking regions of India, such as Delhi and NCR regions, Uttar Pradesh, and Madhya Pradesh. All of them spoke Hindi as their first language, and reported having acquired the language before the age of five. The participants had normal or corrected-to-normal vision and normal hearing, and had no history of any reading disorder or neurological impairment. They were all right-handed, as determined by an abridged version of the Edinburgh Handedness Inventory (Oldfield, 1971). The research protocol for the experiment was approved by the Institutional Ethics Committee (Human) of the Indian Institute of Technology Ropar. The experiment was carried out in accordance with their recommendations. The participants were informed about the entire experimental procedure, and written consent was obtained from them. They were remunerated as per the allowance permitted by the ethical committee. Data from eight further participants were removed from the final analysis either due to excessive EEG artefacts and/or insufficient accuracy in the behavioural task (an error rate of > 25% in any one condition).

### Material

The experimental materials contained 60 sets of sentences in four conditions in the canonical SOV word order (as illustrated in Table 1). The resulting 240 critical sentences were sub-divided and distributed into two lists of 120 sentences each, with each list consisting of 30 sentences per critical condition. Each list containing 120 critical sentences was interspersed with 120 filler sentences of several types. Each participant received one of the two lists, with the order randomized and the list assignment counterbalanced across participants.

Every sentence began with an adverb, followed by the first argument, which was the subject (a masculine, singular, human-animate, common noun). The subject argument was either overtly marked with an ergative case or remained unmarked with the default nominative case. The second argument in the sentence was an unmarked object (always a masculine, singular, inanimate and common noun). Thus, the first and second arguments in the experimental sentences were always animate and inanimate, respectively. Further, all the arguments in the experimental conditions were masculine^2^. The two arguments were followed by the compound light verb structure with two verbs. The first or polar verb was always in the transitive root form, i.e., without any grammatical markers. The second or light verb expressed all required grammatical markers such as agreement, aspect and tense. The light verbs varied in their transitivity (transitive or intransitive verbs) to align themselves with the subject case assignment. A present tense auxiliary concluded the sentence.

For the creation of the 240 critical sentences, five transitive and five intransitive compound light verbs were chosen and repeated over twelve times across the transitive and intransitive conditions^3^. The repetition of the light verb forms was necessary because, in Hindi, only a few verb classes can occur as light verbs, and even fewer are compatible with the required experimental structure involving other entities, such as the NP1 (animate, ergative or nominative case marked), the NP2 (inanimate and default case), the V1(transitive, root form) and the V2 (transitive or intransitive light verb). All experimental sentences consisting of both the critical and filler items and the instructions provided to the participant during the experiment were visually presented in Hindi (in the Devanagari script).

### Procedure

The experiment began with a short practice session followed by the actual experiment, which had 240 sentences divided into six experimental blocks, with the participants taking short breaks between the blocks. First, the participants were informed about the experimental procedure, and a printed instruction sheet explaining the procedure was provided to them. However, the participants were not told the exact question under investigation in the experiment in order to obtain unbiased data. Once they gave informed consent to participate in the experiment, they were required to fill out a questionnaire regarding their linguistic background along with an adapted version of the Edinburgh Handedness Questionnaire (Oldfield, 1971) in Hindi. Following this, the head measurements of the participants were taken, and an appropriately sized Hydrocel GSN net was placed on their scalps. Then, they were seated on a comfortable chair inside a soundproof experimental chamber at a distance of about one meter from the computer screen on which the stimulus was presented using E-Prime v2.0 (Psychology Software Tools, Pittsburgh, PA). Each trial began with a fixation sign “+” at the centre of the screen for a period of 1000 ms, followed by a blank screen for a duration of 200 ms. Rapid serial visual presentation (RSVP) was used for presenting the stimuli in the centre of the computer screen in a chunk by chunk manner (determiner and noun, noun and the case markers, etc., were presented together in a single chunk). Every chunk was presented for a duration of 650 ms with an inter-stimulus interval (ISI) of about 200 ms. This presentation time was selected because of the morphological complexity of the Hindi language and was considered a comfortable reading rate for the Hindi language participants (Schlesewsky et al., 2010; Bhattamishra et al., 2021). Once the fixation sign disappeared, an adverb appeared on the screen, followed by the first argument (NP1), the second argument (NP2), the main verb (V1), the light verb (V2), and finally, the auxiliary (Aux). After this, the participants were required to perform two tasks (Choudhary et al., 2009). First, they performed an acceptability judgment task after the display of the stimulus. As a cue to this task, a “???” sign was displayed for 1500ms on the screen after the auxiliary. Once the participants saw the “???” sign, they had to press a green button in the response pad if they found the just displayed sentence to be grammatically acceptable and, alternatively, a red button in the response pad if they found the sentence to be ungrammatical/unacceptable. Second, after the completion of the acceptability judgement task or after a maximum of 1500 ms duration had passed, the participants performed a probe detection task, whereby a single probe word appeared in the middle of the screen. On seeing the probe word, the participants had to press the green button if they thought that the probe word was present in the experimental stimulus, or, alternatively, the red button if they thought the probe word was not present in the experimental stimulus. The maximum response time allowed for the probe detection task was 3000 ms.

Half of the experimental items were correct sentences, whereas the rest were incorrect sentences. A similar counterbalance was maintained for the probe detection task, with the assignment of an equal number of correct and incorrect probe words. The probe words (correct and incorrect) were constructed in a way that provides a balanced representation of all of the word categories in the experimental stimuli (NP1, NP2, V1, V2). In addition, the distribution of correct or incorrect responses to the left and right response buttons was also counterbalanced. Finally, at the end of the experiment, the participants had to fill out a feedback form, and they were given suitable remuneration for their participation. The entire experimental session, including the electrode preparation, head size measurement, EEG scalp net placement, and stimulus presentation, lasted approximately two hours.

### EEG Recording and data processing

The scalp EEG was recorded using the 33 Ag/AgCl electrodes (32 + VREF) fixed at the scalp with the help of a Hydrocel Geodesic Sensor Net 32 channel (Electrical Geodesics, Philips Neuro, Eugene, Oregon, United States of America). The recordings from the scalp electrodes were referenced to the vertex electrode (Cz) and re-referenced offline to the average activity of the left and right mastoids. The electrooculogram (EOG) was monitosred by means of electrodes placed at the outer canthus of each eye for horizontal eye movements, and above and below the eyes for vertical eye movements. All EEG and EOG channels were amplified using a NET AMPS 400 Amplifier and digitized with a sampling rate of 500 Hz. As per the system recommendations, the inter-electrode impedance was kept below 50kΩ (amplifier input impedance > 1 GOhm) (Ferree, Luu, Russell, and Tucker, 2001).

The EEG data was processed using EEGLAB (Delorme and Makeig, 2004) and the ERPLAB plugin (Lopez-Calderon and Luck, 2014). The raw EEG data was filtered using a bandpass filter of 0.3-30 Hz to remove slow drifts from the signal, followed by offline re-referencing to the average of the two mastoids. Visual inspection was performed to scan the data trials and electrodes, and to remove breaks/pauses between blocks as well as clean the data from any noise or extraordinary artefacts. Following this, automatic rejection of the bad channels was carried out using the ‘pop_rejchan’ function in EEGLAB, whereby data from participants that had >25% bad channels were excluded. This preprocessed original data was filtered again using a bandpass filter of 1-40 Hz for the purpose of computing Independent Component Analysis (ICA). This 1 Hz high-pass filtered data was visually inspected to exclude other kinds of noise, such as non-physiological artefacts like slow drifts arising from electrical devices or wires in/around the recording environment or physiological/biological artefacts such as glossokinetic and other types of body muscle movement (Luck, 2014). Subsequently, ICA was computed, and SASICA plugin was used to detect the artefactual components (Chaumon, Bishop & Busch, 2015). The ICA weights and components marked as artefactual were transferred to the original data, and the artefactual components were rejected from this data. The ERP data were epoched from 200 ms before the onset of the critical word (the light verb) to 1000 ms post-onset to calculate the ERPs for each participant. It was averaged offline for each participant at each electrode within each condition. In the end, a grand average was computed for all the participants (N=30).

### Statistical Data Analysis

The mean acceptability and accuracy, as well as the mean reaction times for the acceptability judgement task and probe detection task, per participant per condition given in Table 1, were calculated for each correct trial using E-Prime 2.0 (Schneider, Eschman, & Zuccolotto, 2002). The statistical analysis of the behavioural data was analyzed by fitting generalized linear mixed-effects models using the lme4 package (Bates et al. 2015) in R (Version 4.4.2, R Core Team, 2024) to examine the factors Case at NP1 (Ergative, Nominative) and the Transitivity of the light verbs at VP2 (Transitive, Intransitive) for the selected participants (N=30).

For the EEG data analysis, the data of a participant was included in the final analyses subject to the following conditions: (1) there should at least be 80% accuracy in the probe detection task, and (2) the data should not contain an excessive amount of EEG artefacts (Kaan and Swaab, 2003). The statistical analysis was performed by means of linear mixed-effects models using the lme4 package in R to examine the factors Case (Ergative, Nominative) and Transitivity (Transitive, Intransitive) and the topographical factor Region of Interest (ROI). Based on visual inspection of the data and previous literature, mean amplitude values for each condition were extracted for time windows of interest (Choudhary et al., 2009, Bornkessel-Schlesewsky, Kretzschmar, Tune, Wang, Genc, Philipp, Schlesewsky, 2011). Furthermore, for the purpose of statistical analysis, selected groups of electrodes were clustered and averaged together to form four lateral and six midline ROIs. Figure 1 illustrates the lateral and midline sites, which underwent separate statistical analyses (Bornkessel-Schlesewsky et al., 2011). The three lateral regions were as follows: left-anterior (3, 5, 11, 13), left-posterior (7, 9, 15, 25), right-anterior (4, 6, 12, 14), and right-posterior (8, 10, 16, 26). For the midline region, each electrode was analyzed as a ROI of its own: 18, 27, 17, 28, 19, 20.

**Figure 1:**
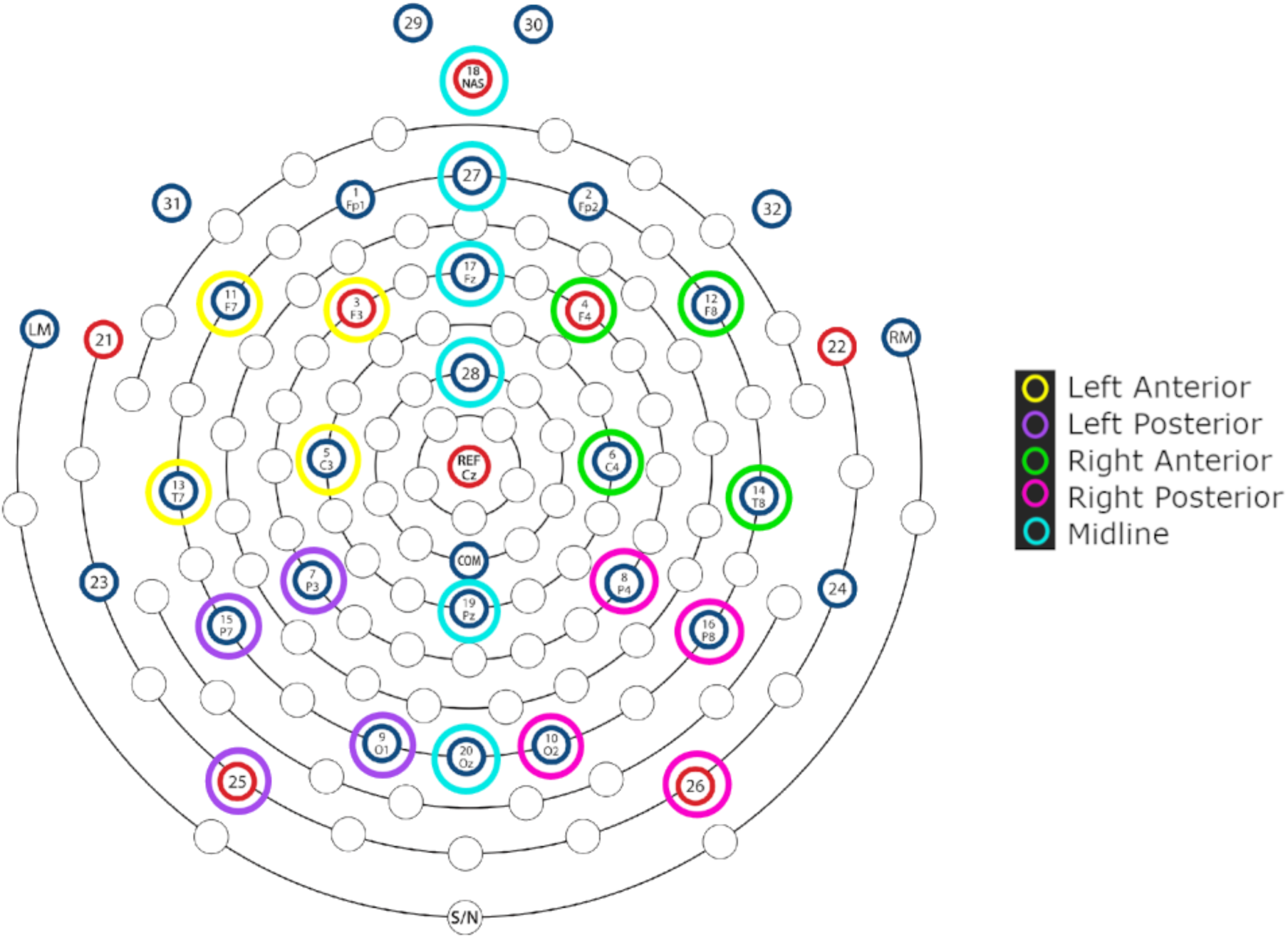
Electrode grouping of selected regions of interest for statistical analysis (Adapted from Hydrocel Geodesic Sensor Net 32 Channel montage by Magstim, EGI)

## Result

### Behavioural Results

Table 1 shows the mean acceptability judgement and mean probe accuracy for the critical conditions, illustrating that there was higher acceptability for grammatical conditions than ungrammatical ones. The acceptability judgements are presented in the form of a raincloud plot (Allen, Poggiali, Whitaker, Marshall, and Kievit, 2019) in Figure 2. Figure 2A shows the by-participant variability of acceptability ratings, with the individual data points representing the mean by-participant acceptability of each Case and Transitivity combination. Figure 2B shows the by-item variability of acceptability ratings, with the individual data points representing the mean by-item acceptability of each Case and Transitivity combination. The relatively low acceptability for the ergative violation condition (*EI) compared to the nominative violation condition (*NT) is in line with the theoretical descriptions of the syntactic structures in the Hindi language. That is, the ergative case assignment is a non-default rule in the language and it occurs only in a strict structural pattern, unlike the nominative case assignment which occurs as a default rule in the language, and therefore, an ergative case violation is less likely to be accepted by Hindi native speakers. In the probe detection task, the accuracy of the participants was at ceiling level across all four conditions.

**Figure 2.**
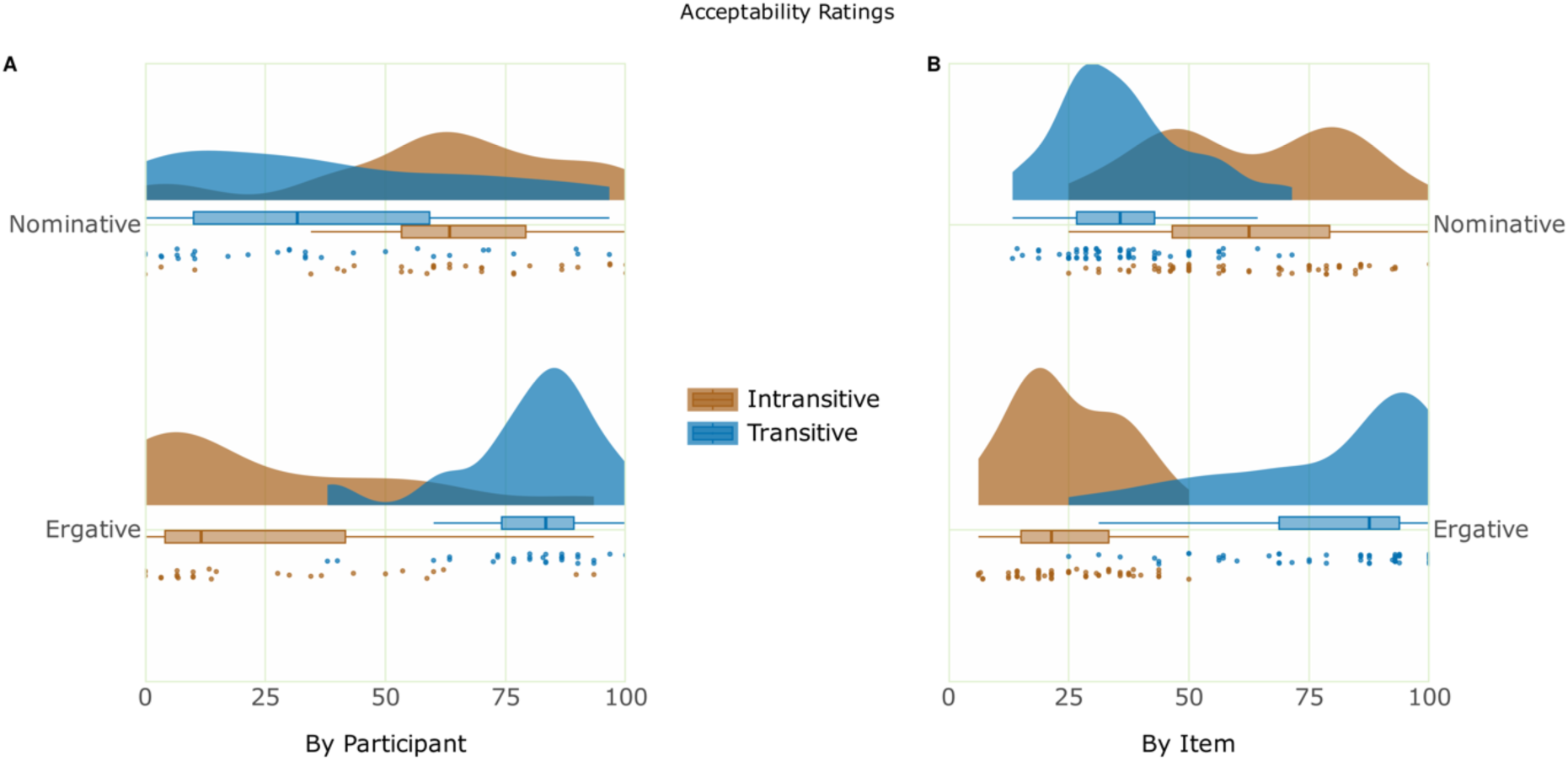
Raincloud plot of the behavioural data. Panel A depicts the mean acceptability ratings by Participant. Panel B depicts the mean acceptability ratings by Item.

The behavioural data was analyzed by fitting generalized linear mixed-effects models using the lme4 package in R to examine the relationship of sentence acceptability and accuracy to the factors Case and Transitivity. Categorical factors used scaled sum contrasts (i.e., coefficients reflect differences to the grand mean), which “makes the estimated slopes the same as for treatment coding and easier to interpret” (Schad et al., 2020, p.7). Model selection was based on the Akaike Information Criterion (AIC), whereby the model with the lowest AIC was selected as the one that best explained the data. The analysis of the acceptability data showed that the model involving the factors Case and Transitivity, and their interaction term, with random intercepts for participants and items, and by-participant and by-item random slopes for the effect of Case and Transitivity, best explained the acceptability data (AIC = 3289.2). Models involving less complex / no random slopes specifications either did not converge or had a higher AIC. Type II Wald chisquare tests on the selected model showed a main effect of Transitivity (χ2(1) = 5.98, p = 0.01) and an effect of the interaction Case x Transitivity (χ2(1) = 618.57, p < 0.001). Estimated marginal means on the response scale were computed on the model using the emmeans package (Lenth, 2021) to resolve this interaction, which showed that the pairwise contrasts between estimates of Intransitive vs. Transitive light verbs within each level of Case showed a simple effect of Transitivity both when the Case was Ergative (estimate = 0.704, SE = 0.0339, p < 0.001) as well as when it was Nominative (estimate = -0.365, SE = 0.0502, p < 0.001). The analysis of the answering accuracy showed no effects. The reaction time of the two tasks was not further analyzed as the tasks were not directly time-locked to the critical region of the sentence but instead were meant for the whole sentence (similar pattern as in Choudhary et al., 2009).

### ERP Results

Visual inspection and statistical analyses revealed two ERP effects at the critical position of the light verb: a negativity for the ergative case + intransitive light verb violation condition in the time range of 350–550 ms shown in Figure 3 (Figure 3a and 3b) and a late positivity, which was broadly distributed in the time range of 750–950 ms for the nominative case + transitive light verb violation condition, as shown in Figure 4 (Figure 4a and 4b). The ERP data were statistically analyzed by fitting mixed effects models using the lmer function from lme4 package in R, with fixed effects Case, Transitivity, ROI and their interaction terms. Categorical factors used scaled sum contrasts (i.e., coefficients reflect differences to the grand mean). Instead of performing a traditional baseline correction, the mean prestimulus EEG amplitude (-200 to 0 ms) was included in the model as a scaled and centred covariate (Alday, 2019). However, we do not interpret effects involving prestimulus amplitudes, in line with the fact that these did not form part of our hypotheses, but also because they were included in the model to account for and regress out potential contributions of the baseline period to the critical period. It is indeed valid to “include additional covariates as controls without further interpreting those covariates” (Alday 2019, p.9). Instead, we focus on effects attributable to the experimental factors Case and Transitivity, and for each model interpret the highest order interaction involving both of these factors. Additional interactions with prestimulus interval are nevertheless resolved for a range of prestimulus amplitude values from −5 to 5 μV, which showed that the overall pattern of results remained broadly consistent (see supplementary supporting information online).

**Figure 3.**
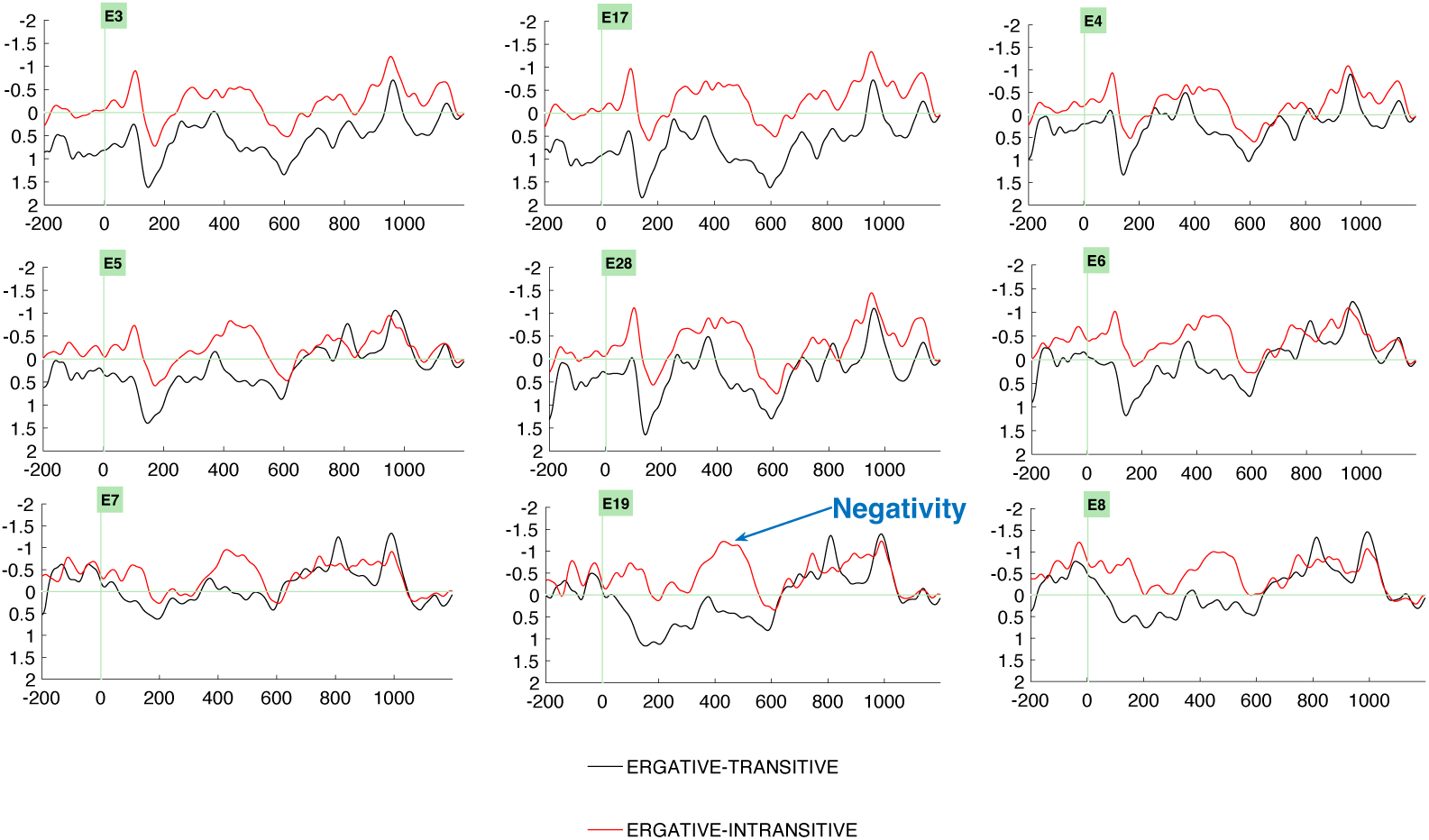
(a) Grand average ERPs (N=30) at the light verb position comparing the control condition, Ergative-Transitive (marked in black), and the critical-violation condition, Ergative Intransitive (marked in red), reveals a negativity effect for the Ergative Intransitive conditions. Negativity is plotted upwards by convention. Abbreviation: Left Anterior electrodes (3, 5), Midline electrodes (17, 28, 19), Right Anterior (4, 6), Left Posterior electrodes (7), Right Posterior electrodes (8).

**Figure 3.**
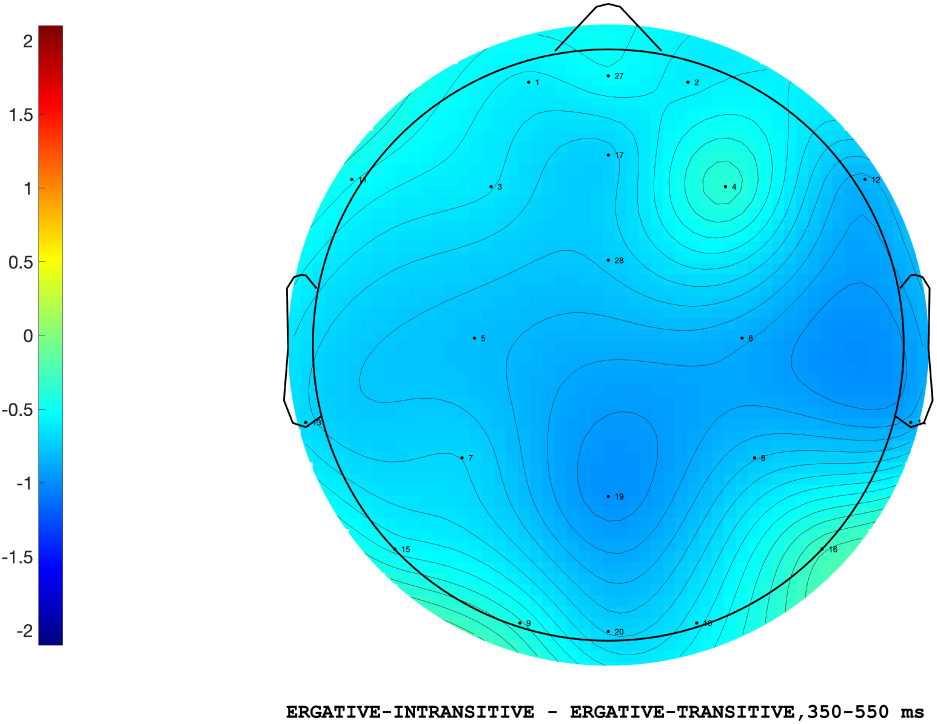
(b) Topographic difference between the ERP amplitudes in the ergative ungrammatical (*EI) and grammatical (ET) conditions from the position of the light verb in the time window 350-550ms.

**Figure 4.**
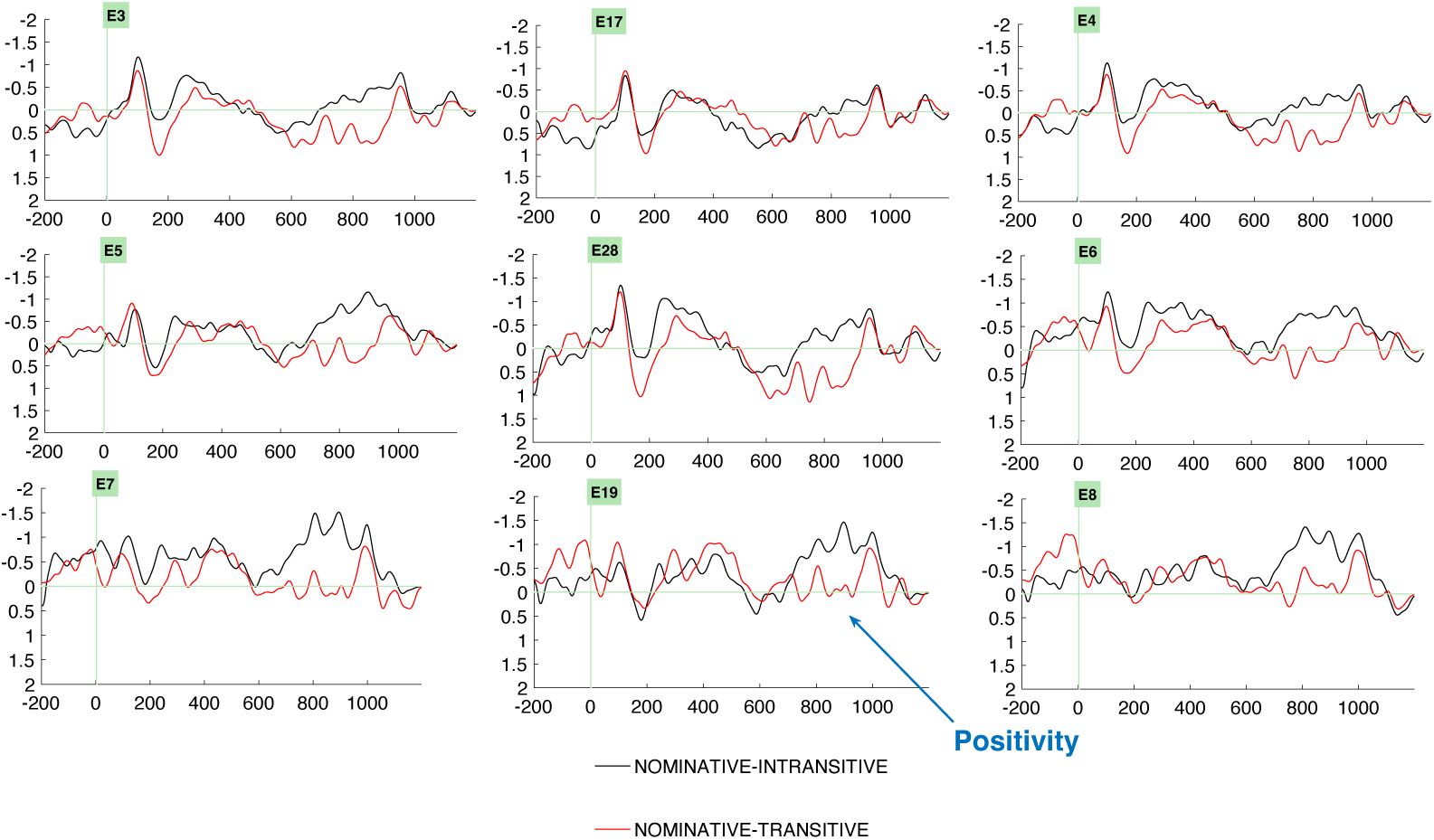
(a) Grand average ERPs (N=30) at the light verb position comparing the control condition, Nominative-intransitive (marked in black), and the critical-violation condition, Nominative Transitive (marked in red), reveals a positivity effect for the Nominative transitive conditions. By convention, negativity is plotted upwards. Abbreviation: Left Anterior electrodes (3, 5), Midline electrodes (17, 28, 19), Right Anterior (4, 6), Left Posterior electrodes (7), Right Posterior electrodes (8).

**Figure 4.**
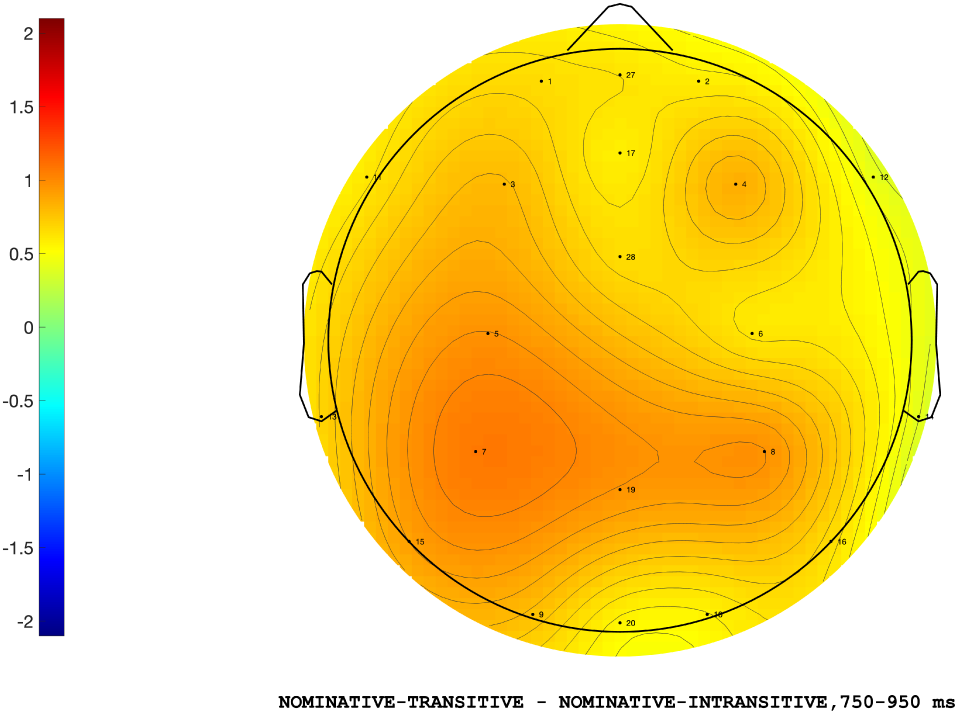
(b) Topographic difference between the ERP amplitudes in the nominative ungrammatical (*NT) and grammatical (NI) conditions from the position of the light verb in the time window 750-950ms.

### 350–550 ms time-window

The analysis in the lateral regions of interest showed that the model involving the factors ROI, Case and Transitivity, and their interaction terms, with random intercepts for participants, and by-participant random slopes for the effect of Case, Transitivity and their interaction term best explained the mean ERP amplitude (cAIC = 1329.35). Models involving less complex / no random slopes specifications either did not converge or had a higher cAIC. Type II Wald chisquare tests on the selected model showed an effect of the interaction Case x Transitivity (χ2(1) = 4.28, p = 0.03). Estimated marginal means on the response scale were computed on the model using the emmeans package (Lenth, 2021) to resolve this interaction, which revealed that the pairwise contrasts between estimates of Intransitive vs. Transitive light verbs within each level of Case showed a simple effect of Transitivity when Case was Ergative (estimate = 0.5758, SE = 0.234, p = 0.01).

The analysis in the midline regions of interest showed that the model involving the factors ROI, Case and Transitivity, and their interaction terms, with random intercepts for participants, and by-participant random slopes for the effect of Case, Transitivity and their interaction term best explained the mean ERP amplitude (cAIC = 2569.05). Models involving less complex / no random slopes specifications either did not converge or had a higher cAIC. Type II Wald chisquare tests on the selected model showed an effect of the interaction Case x Transitivity (χ2(1) = 7.17, p = 0.007). Estimated marginal means on the response scale were computed on the model to resolve this interaction, which revealed that the pairwise contrasts between estimates of Intransitive vs. Transitive light verbs within each level of Case showed a simple effect of Transitivity when Case was Ergative (estimate = 0.7151, SE = 0.284, p = 0.01).

### 750–950 ms time-window

The analysis in the lateral regions of interest showed that the model involving the factors ROI, Case and Transitivity, and their interaction terms, with random intercepts for participants, and by-participant random slopes for the effect of Case, Transitivity and their interaction term best explained the mean ERP amplitude (cAIC = 1325.39). Models involving less complex / no random slopes specifications either did not converge or had a higher cAIC. Type II Wald chisquare tests on the selected model showed an effect of the interaction Case x Transitivity (χ2(1) = 3.83, p = 0.05). Estimated marginal means on the response scale were computed on the model to resolve this interaction, which revealed that the pairwise contrasts between estimates of Intransitive vs. Transitive light verbs within each level of Case showed a simple effect of Transitivity when Case was Nominative (estimate = 0.457, SE = 0.227, p = 0.05).

The analysis in the midline regions of interest showed that the model involving the factors ROI, Case and transitivity, and their interaction terms, with random intercepts for participants, and by-participant random slopes for the effect of Case, and their interaction term best explained the mean ERP amplitude (cAIC = 2718.99). Models involving less complex / no random slopes specifications either did not converge or had a higher cAIC. Type II Wald chisquare tests on the selected model showed a marginal effect of the interaction between ROI x Case x Transitivity (χ2(5) = 10.33, p = 0.06). Estimated marginal means on the response scale were computed on the model to resolve this interaction, which revealed that the pairwise contrasts between estimates of Intransitive vs. Transitive light verbs within each level of ROI and Case showed no effect of Transitivity.

## Discussion

The current study investigated the processing of ergative case in a split-ergative language to examine whether the underlying comprehension mechanism behind ergative case processing is similar to that of nominative case, and whether the neurophysiological correlates are in line with previous studies on ergative languages. Moreover, unlike the existing studies in the literature, the current study used light verbs to examine if structural variation plays a role in the processing of ergative case.

At the critical position of the light verbs, the results revealed qualitatively different electrophysiological correlates for the processing of ergative and nominative case-based violations. The ergative case violation condition was anomalous due to the incongruous combination of an ergative case-marked animate subject noun with an intransitive light verb form. This evoked a centroparietal negativity effect between 350 and 550ms in contrast to its grammatical counterparts. This effect could be plausibly interpreted as an instance of an N400 effect in view of its topographical distribution and peak latency. By contrast, the nominative violation condition had a nominative case assignment rule mismatch due to the presence of a transitive form of the light verb. A broadly distributed late positivity effect between 750 and 950ms was observed for the nominative violation condition, which could be interpreted as a P600 effect.

We formulated two competing hypotheses regarding the neurophysiological correlates of case violations in Hindi. Our primary hypothesis, grounded on the notion of similar correlates for case processing mechanisms across languages, predicted no difference in the neural responses to ergative and nominative case violations. In contrast, our alternative hypothesis, informed by previous research (Choudhary et al., 2009), posited that the processing correlates of ergative and nominative cases should differ qualitatively, and that this would be observable as ERP differences at the position of light verbs. Specifically, we anticipated that, in line with the previous study by Choudhary et al. (2009), nominative case violations would elicit an N400 effect, whereas ergative case violations would evoke a biphasic N400-P600 effect.

Our results provide strong evidence for the ergative case being processed differently, suggesting that the predictions that the two kinds of subject cases (nominative and ergative) hold for sentence comprehension, as well as the structural possibilities that they entail, are quite distinct. However, our results differ from the previous findings in Hindi (Choudhary et al., 2009: N400-P600 for ergative case violation and N400 for nominative case violations; Dillon et al., 2012: RAN-P600 for ergative case violation) as well as other ergative languages such as Punjabi (Gulati, 2021: early positivity for nominative case violations and another positivity for the ergative case violations), and Basque (Zawiszewski et al., 2011: N400-P600 for double ergative case violation; Díaz et al., 2011: P600 for ergative case violation).

Considering the findings from previous research (Choudhary et al., 2009; Dillon et al., 2012; Gulati, 2021; Díaz et al., 2011; Zawiszweski et al., 2011) in conjunction with results from the present study, it is clear that multiple factors contribute to variability in ERP studies on case processing. A closer examination reveals that in ergative case processing studies, it is crucial to distinguish between two types of violations: those that arise from the case-based relationship between the verb and its arguments, and those that occur when arguments are analyzed independently of the verb. The location of the case violation, the complexity of the verbal structures and the sentence types, and the word order (canonical or non-canonical) are all important factors in understanding the variations in case processing. In this regard, it is noteworthy that previous studies from within the same language, Hindi (Choudhary et al., 2009; Dillon et al., 2012) and from a typologically similar language, Punjabi (Gulati, 2021), have explored only one facet of split-ergativity. Specifically, these studies have shown that in Hindi, split-ergativity is conditioned by the verb’s grammatical aspect and/or tense information. Consequently, a change in aspect type can lead to variations in the processing of ergative case, highlighting the complex interplay between these linguistic factors. The current study examined a hitherto unexplored facet in ergative case processing studies, namely, the interplay of transitivity of light verbs and subject case assignment. Thus, by keeping the tense and aspect information constant, with solely a transitivity variation being realized in a compound light verb construction, stimuli in the current study ensured that no other morphosyntactic manipulation occurred at any position.

Complex predicate structures such as compound light verb constructions -- in which it is often not so easy to define the prototypical transitivity of the sentence due to its intricate mapping of semantic and syntactic transitivity onto two verbs -- are quite different from a simple transitive sentence with a single verb (Butt, 1995). The different kinds of sentential structures can change the way we understand and represent events. When ergativity is processed in the context of a compound light verb construction, the transitivity of the light verbs is of paramount importance since it determines the subject case assignment in Hindi. Our results confirm that in complex structures such as the ones used in our study, the transitivity of the light verbs indeed modulates the processing of ergative case. The current study thus adds another dimension to our understanding of case processing, which could not have been investigated using simple transitive structures as in previous studies. As evidenced by our results, variation in sentence structure can significantly influence case processing, particularly ergative case processing. This emphasizes the importance of considering language-specific characteristics when examining a linguistic phenomenon. In the following sections, we discuss the functional interpretation of the N400 and P600 effects evoked in the study and outline our propositions for future research.

### The N400 Effect

Classically, the N400 component is said to be reflective of lexical-semantic processing (Kutas and Hillyard, 1984; Federmeier and Kutas, 1999; Kutas and Federmeier, 2000). However, further research suggests that it is also associated with issues arising from syntactic/morphological processing. Thus, N400 effects have been reported for revision or reanalysis of case marker and related predictions due to thematic interpretative conflict (Friederici and Frisch, 2000; Frisch and Schlesewsky, 2001, 2005; Mueller Hirotani and Friederici, 2007), word order-based and grammatical relation reanalysis (Hopf, Bayer, Bader and Meng, 1998; Bornkessel et al., 2004, Haupt et al., 2008) and other kinds of reanalysis induced by revisions of case marking (Bornkessel et al., 2003, 2004; Coulson et al., 1998; Rösler, Pütz, Friederici, and Hahne Pechmann, 1993; Schlesewsky and Bornkessel, 2004; Schlesewsky, Bornkessel, and Frisch, 2003). Some previous studies have interpreted the N400 effect as an indicator of difficulty in thematically hierarchizing the arguments or problems in assigning thematic roles to the noun phrase arguments (Choudhary, 2011; Frenzel, Schlesewsky and Bornkessel-Schlesewsky, 2011) as well as for linguistic cue-based conflicts (Roehm et al., 2004; Philipp, Bornkessel-Schlesewsky, Bisang, and Schlesewsky, 2008). The N400 effect is also said to be reflective of discourse integration costs (Burkhardt and Roehm, 2007) as well as case and aspect-related violations that cause a misalignment of sentential interpretations (Bornkessel-Schlesewsky and Schlesewsky, 2009; Choudhary et al., 2009). Specifically in the context of studies on ergative languages, Choudhary et al. (2009) examined the processing of nominative and ergative case in Hindi, and reported an N400 effect for case violations, which they interpreted as resulting from a mismatch of an interpretively relevant rule. Zawiszewski et al. (2011) reported an N400 effect for ergative case violations in Basque, with the effect said to reflect the difficulty in ascribing a thematic function to an ungrammatical subject case marked argument.

The N400 observed in the present study can be interpreted in line with many of the interpretations provided in previous studies. However, we suggest that the effect observed in our study more precisely aligns with that of the Choudhary et al. (2009), who interpreted the N400 effect to be reflective of an interpretively relevant rule mismatch. They observed an N400 effect for both ergative and nominative case violations, with the amplitude of the effect modulated by the case that engendered the violation. That is, the effect was larger for ergative case violations than nominative case violations, which they attributed to the interpretive rule mismatch being more severe for ergative case violations. In the ergative violation condition in our study, the anomaly at the intransitive light verb causes a severe breakdown of the previously constructed interpretation. The breakdown of the processing mechanism can be explained as follows: In languages exhibiting a split-ergative pattern, when the processing mechanism encounters an ergative-marked subject argument, it generates a prediction that the subsequent utterance will be transitive, featuring a simple transitive verb in the perfective aspect. This interpretation does not change when the processor encounters the object argument in our stimuli. However, upon encountering the polar verb V1, the processor revises its initial prediction, recognizing that the predicate structure is actually a compound light verb construction rather than a simple transitive verb. As a result, it updates its expectation to anticipate a transitive light verb. In the control ergative condition, the processor encounters a transitive light verb, which can assign an ergative case to the subject argument, and thus, the processing mechanism does not encounter a problem. However, in the ergative violation condition, instead of a transitive light verb, the parser encounters an intransitive light verb, which leads to a breakdown of interpretation as the intransitive light verb cannot assign ergative case to its subject argument, leading to the elicitation of the N400. Further, unlike the combination of a transitive polar and a transitive light verb in the control condition, which makes the syntactic structure transitive, the intransitive light verb in the violation condition causes a reduction in the syntactic transitivity of the sentence. This is because of its failure to assign ergative case to the subject of the sentence and, consequently, leads to an ill-formed sentence. If the presence of an ergative argument is a strong indicator of transitivity and/or agency/volition (Hook, 1974; Butt, 1993; Mohanan, 1994), then the intransitivity of the light verb would entail a reduced agency/volition on the part of the agent argument and, in turn, an overall reduction in the syntactic transitivity of the sentence. This would suggest that the presence of an object and/or the assignment of the ergative case can serve as indicators for establishing the semantic transitivity of a sentence. Despite this, semantic transitivity does not always entail syntactic transitivity, or syntactic transitivity does not necessarily trigger the transitive meaning (Drocco, 2018; Drocco and Tiwari, 2020). We interpret the absence of a positivity in this condition as a consequence of the lack of an alternative structure, which could have initiated the process of repair and reanalysis. In Hindi, an initial argument marked with an ergative case, as in the current study, offers a restrictive structural environment for constructing the sentence meaning. The parser only has one sentential interpretation from the very first argument, which it maintains until the very last element (light verb) is encountered. Therefore, there is no possibility of reanalyzing into alternative structural configurations. Moreover, the current study had no additional instances of morpho-syntactic or semantic discrepancies at any other position in the violation condition. Thus, there was no alternative interpretation possible.

### The P600 Effect

In the sentence processing literature, the late positivity or the P600 effect, also sometimes called the syntactic positive shift, has been initially interpreted as a marker for syntactic/structural processing and syntactic ambiguities (Osterhout and Holcomb, 1992; Hagoort, Brown and Osterhout, 1999; Hagoort, Brown, and Groothusen, 1993; Hagoort and Brown, 2000). The late positivity is often associated with reanalysis of the morpho-syntactic structure (Friederici, 1995, 2002), especially diagnostic and structural reanalysis (Friederici, 1998; Fodor and Inoue, 1994) as well as any kind of linguistic repair and revision processes whenever a structural integration difficulty or complexity in syntactic analysis arises (Kaan, Haris, Gibson and Holcomb 2000; Friederici, Hahne and Saddy, 2002; Kaan and Swaab, 2003). It has also been linked with processing all kinds of linguistic/grammatical misfit words or errors (Coulson, King and Kutas, 1998; Frisch and Schlesewsky, 2001, 2005; Mueller, Hahne, Fujii, and Friederici, 2005; Mueller, Hirotani and Friederici, 2007; Rossi, Gugler, Hahne and Friederici, 2005; Nevins, Dillon, Malhotra, and Phillips, 2007). The P600 is also considered an index of well-formedness/ill-formedness assessment (Bornkessel and Schlesewsky, 2006) and is further said to be reflective of the general conflict monitoring process, particularly when induced by inconsistencies between different information types, such as syntactic and thematic/plausibility information (Frenzel, Schlesewsky, Bornkessel-Schlesewsky, 2011; van Herten, Chwilla and Kolk, 2006; Bornkessel-Schlesewsky and Schlesewsky, 2008). It has also been observed in sentences that are grammatically correct but semantically anomalous or implausible, namely semantic reversal anomalies (Kim and Osterhout, 2005; van Herten, Kolk, and Chwilla, 2005, and van Herten, Chwilla and Kolk, 2006), and more generally for semantic integration problems (Brouwer, Fitz, Hoeks, 2012; Brouwer, Crocker, Venhuizen, Hoeks, 2017).

In previous studies on ergative languages, Choudhary et al., (2009) in Hindi reported a positivity effect for the ergative case violation, which they interpreted as a marker for the over-application of a non-default morphosyntactic case rule which detects the incompatibility between the subject case and aspect marker. In another study in Hindi, Dillion et al., (2012) also reported a late positivity for syntactic (ergative case-induced) and semantic (adverb-induced) violations, which was interpreted as a marker showing the presence of linguistic errors. However, since the predictions made by the syntactic cues (ergative case) were considered more robust, the P600 effect was also more prominent in that condition. The study by Dillon et al. (2012) shows how a P600 effect can also be evoked for case-induced linguistic errors. In Punjabi, Gulati (2021) found late positive effects of differing latencies for the processing of ergative and nominative cases and verbal aspect-based violations, which were attributed to an ill-formedness of the sentence (Bornkessel and Schlesewsky, 2006, 2008). In Basque, both Díaz et al., (2011) and Zawiszewski et al., (2011) reported positivity effects for ergative case violation sentences. While the former interpreted it as a marker indicating conflict monitoring mechanisms, the latter suggested it to be a correlate of reanalysis/repair and identification of incorrectly formed case-based sentences.

In the present study, there are potentially several sentence structures and interpretations possible after encountering an initial nominative human argument. Based on its animacy, as suggested by the subject/agent first theory, the first argument can be assumed to be the subject of the sentence (Bickel et al., 2015), and the sentence can be interpreted as having an intransitive reading based on the minimality hypothesis (Chomsky, 1993, 1995). However, this prediction is later abandoned on encountering the second argument, which being an inanimate entity, is processed as the object argument. At the second argument, the parser then assumes a transitive reading of the sentence, which is again dropped when the polar verb is encountered. A transitive polar verb is indicative of a complex predicate structure. Even at this position, there are numerous other viable structural interpretations. Thus, when a transitive light verb in the perfective aspect is encountered in the violation conditions, the light verb cannot validate the assignment of nominative case to the subject because a light verb with such a specification can only assign ergative case in Hindi. This results in an unintended increase in the syntactic transitivity of the complex predicate structure, which is not required for the assignment of nominative case. Despite the mismatched transitivity of the light verb causing a nominative case violation at the light verb position, this violation lacks the potential to result in an interpretatively relevant case-based rule violation as severe as the one in the compound light verb constructions with ergative case violations. This is because even when the parser’s previously chosen most optimal nominative case-based structural interpretation fails, it does not entirely hinder the sentence comprehension process. Throughout the parsing stage, the parser has other potential structural configurations active. As a result, even if one of the interpretations fails, the parser still has access to alternative interpretations. By selecting the subsequent best interpretation, the parser possibly reanalyzes and fixes the structure, leading to the late positivity effect. Hence, we interpret the late positivity effect in our study as a marker for reanalysis, which has consistently been associated with a P600 effect in the literature (Friederici, 1995, 1998, 2002; Friederici, Hahne and Saddy, 2002; Kaan, Harris, Gibson and Holcomb, 2000; Kaan and Swaab, 2003).

Summing up, the results of the current study reveal processing differences between ergative and nominative cases in Hindi, and provide robust evidence for the differential processing of ergative case violations in Hindi light verb structures. Ergative case violations in a complex predicate structure and the illicit use of an intransitive light verb resulted in an N400 effect, while the illicit use of a transitive light verb in the nominative violation condition resulted in a P600 effect. This highlights the importance of the transitivity of light verbs in the processing of case violations in Hindi. Moreover, a comparison between the current and previous studies in ergative languages (Basque, Punjabi, Hindi) would reveal that, unlike those studies which used simple transitive structures, the current study employed light verbs, whereby the transitivity of the light verb controls the case assignment. The fact that there were qualitative differences in the neurophysiological correlates observed between the current and existing studies points towards transitivity and structural variation being possible causes for the neurophysiological variations both within and across languages. Taken together, these findings foreground the need for probing into language-specific properties and taking structural variations into account rather than looking for universal processing strategies for a particular linguistic phenomenon across languages of the world. We suggest that to understand the full extent of the influence of ergativity and to decipher the role that case in general plays as a processing cue in language comprehension, more cross-linguistic work from understudied languages is required. Thus, focusing on structure-specific features by incorporating diverse syntactic structures and understanding the language-specific properties that contribute to ergative case processing are important points to consider for future research.

## Conflict of Interest

The authors declare that the research was conducted in the absence of any commercial or financial relationships that could be construed as a potential conflict of interest.

## Author Contributions

AMM: Conceptualization, Investigation, Methodology, Project administration, Resources, Software, Data curation, Formal analysis, Validation, Visualization, Writing – original draft, Writing – review & editing. RM: Software, Data curation, Formal analysis, Funding acquisition, Supervision, Validation, Visualization, Writing – review & editing. MG: Validation, Writing – review & editing. SB: Supervision, Validation, Writing – review & editing. KKC: Conceptualization, Methodology, Funding acquisition, Supervision, Validation, Visualization, Writing – review & editing.

## Funding

This work was supported by the Indian Institute of Technology Ropar through an ISIRD grant Ref no. IITRPR/Acad./2216.

## Acknowledgements

The work was supported by the Department of Science and Technology, Govt. of India, through a project to Dr. Kamal Kumar Choudhary (SR/CSI/141/2012). We would like to acknowledge the Indian Institute of Technology Ropar (IIT Ropar) for the research facility maintained through an ISIRD grant to Dr Kamal Kumar Choudhary [IITRPR/Acad./2216].

## Data Availability Statement

The datasets generated/analyzed for this study can be found in the following repository: https://doi.org/10.5281/zenodo.14975073

## Footnotes

1 The compound light verb constructions fall under the umbrella term of ‘complex predicate structures’ and have a verbal structure where two or more predicational elements (nouns, verbs, and adjectives that can usually stand entirely on their own) combine to form a single, but a complex event, such that their biclausal arguments map onto a monocausal syntactic structure (Butt, 1993b; Butt and Lahiri, 2013). There is an enormous amount of typological literature available on complex predicate structures, their exact definition, the subdivision within, and the theoretical stance taken by researchers on them, especially in South Asian languages (Kachru, 1978; Hook, 1974, 1991; Butt, 1995; Butt and Lahiri, 2002). However, there’s a dearth of studies on the experimental manifestation of their mental representation. An in-depth understanding of how these structures are processed becomes necessary to account for differences in the morphosyntactic and semantic constraints on potential V1-V2 combinations and the consequential effect on the sentence structure (detailed review of the descriptive and typological work on different kinds of Hindi complex predicates structures: Hook, 1974, 1977, 1988, 1991; Nespital, 1966, 1989, 1997; Kachru, 1993, 2006; Verma, 1993; Butt, 1993b, 1995, 1997, 2006, 2010; Butt and Ramchand, 2001; Masica, 1976; Kachru and Pandharipande, 1980; Abbi and Gopalakrishnan, 1991; Hook and Pardeshi, 2005; Poornima, Jean-Pierre, Koenig 2008, 2009; Poornima, 2012; Slade, 2013).

2 All the arguments used in the critical sentences were in the masculine form because the feminine arguments have a completely different agreement alignment pattern in the Hindi language. The usage of feminine arguments would cause further alteration in the compound light verb structures as the light verb would now be responsible for an interplay of both case and agreement assignment. Hence, to maintain the structural manipulation solely restricted to case-based variations, we used exclusively masculine nouns in the experiment. acceptability in the acceptability judgement task; RT1: reaction time for the acceptability judgement task; Accuracy: mean accuracy in the probe detection task; RT2: reaction time for the probe detection task.

3 The selection criteria for transitive and intransitive light verbs in Hindi were based on the most frequently occurring light/vector verbs from the Hindi/Urdu Treebank (Palmer, Bhatt, Narasimhan, Rambow, Sharma, and Xia, 2009; Vaidya, Agarwal, Palmer, 2016) and based on the inclusion and exclusion assessment for light verbs (Bahl, 1964; Kachru, 1966; Pray, 1970; for classification of light verbs in relation to each other, refer to Hook, 1991). Some typological studies have also provided diagnostic measures for evaluating Hindi light verb constructions, although they may not be pertinent to every scenario and primarily adhere to transitive light verbs and conjunct light verb constructions (Mohanan,1994; Butt, 2010; Butt and Lahiri, 2013; Bhattacharyya, Chakrabarti, Sarma, 2007).

## Notes

### Competing Interest Statement

The authors have declared no competing interest.

